# The Antagonism of Folate Receptor by the Integrase Inhibitor Dolutegravir: Developmental Toxicity Reduction by Supplemental Folic Acid

**DOI:** 10.1101/576272

**Authors:** Robert M. Cabrera, Jaclyn P. Souder, John W. Steele, Lythou Yeo, Gabriel Tukeman, Daniel A. Gorelick, Richard H. Finnell

## Abstract

Human immunodeficiency virus (HIV) integrase inhibitors are increasingly being used for antiretroviral therapy (ART), and dolutegravir (DTG/Tivicay) has emerged as a leading core agent. In 2018, the Tsepamo study reported a 6- to 9-fold increase for neural tube defect (NTD) risk among the offspring of mothers receiving DTG during early gestation. Maternal folate (vitamin B9) status is the largest known modifier of NTD risk, so we evaluated folate-related mechanisms of action and the critical period for DTG developmental toxicity. Folate receptor (FOLR1) binding studies indicate DTG is a non-competitive FOLR1 antagonist at therapeutic concentrations. *In vitro* testing indicates calcium (2mM) increases FOLR1-folate interactions and alters DTG-FOLR1-folate interactions and cytotoxicity. DTG does not inhibit downstream folate metabolism by dihydrofolate reductase (DHFR). Early embryonic exposure to DTG is developmentally toxic in zebrafish, and supplemental folic acid can mitigate DTG developmental toxicity. The results from these studies are expected to inform and guide future animal models and clinical studies of DTG-based ART in women of childbearing age.

---

The human immunodeficiency virus and acquired immune deficiency syndrome (HIV/AIDS) epidemic remains a prominent public health challenge. Globally, there are approximately 36.7 million people living with HIV, with 2.1 million new infections occurring annually, resulting in 1.1 million AIDS-related deaths per year^1^. Current HIV antiretroviral therapy (ART) includes C-C chemokine receptor type 5 (CCR5) antagonists, fusion inhibitors, integrase inhibitors, nucleoside and non-nucleoside reverse transcriptase inhibitors (NRTIs and NNRTIs), and protease inhibitors. Dolutegravir (DTG) is the clinically preferred integrase inhibitor used alone (Tivicay, ViiV Healthcare), or in combination (e.g. Triumeq, ViiV Healthcare) as ART for HIV infection in adults and children. A teratogenic risk for DTG was recently reported after neural tube defects (NTDs) were observed in four infants from mothers who had been taking DTG at the time of conception in Botswana^2^. The World Health Organization (WHO) subsequently provided guidelines that DTG use be avoided by women of childbearing potential unless they used adequate contraception methods^3^. Based on this cohort study, the current guidelines are warranted and prudent, but more evidence is needed before scientific and medical communities can support a causal relationship between DTG and developmental toxicity, including NTDs.

## Developmental Toxicity of DTG

There is increasing usage of the HIV integrase inhibitors are for ART, and DTG has emerged as the leading compound in this class of medications. The DTG/Tivicay manufacturer reports (US FDA approved Tivicay Prescribing Information, 09/2018) animal reproduction studies showed no evidence of adverse developmental outcomes. Specifically, DTG was administered orally at up to 1,000 mg per kg daily, in rats and rabbits, during the period of organogenesis, days 6-17 and 6-18, respectively. Although not statistically significant, examination of these data indicates one occurrence (1/167) of cranioschisis (anterior NTD) at 40 mg per kg in rabbits (Oral Study for Effects of S-349572 Sodium on Embryofetal Development in Rabbits). An ongoing observational human cohort study in Botswana reported a 6-to 9-fold increase for NTD risk in the offspring from mothers receiving DTG^2,4^.

## Neural Tube Defects and Folate

NTDs occur when the development of the neural tube, the primordial central nervous system that gives rise to the of the brain and spinal cord, is damaged or impaired during the critical period (human gestational days 17-30) during embryonic development. The largest reported modifier for NTD risk is folate (vitamin B9) status, so pregnant women and women of child bearing age are advised by the World Health Organization (WHO) and the US Public Health Service (USPHS) to take supplemental folic acid (FA) to ensure a daily intake of 400μg^5-8^. Folic acid is a water-soluble provitamin that is transformed to bioactive forms of folate *in vivo* ^9^. Reduced forms of folate such as Leucovorin, also known as 5-formyltetrahydrofolate (folinic acid, SFA), or 5-methyltetrahydrofolate (Metafolin, 5-CH_3_-THF) are active forms and do not require enzymatic reduction by dihydrofolate reductase (DHFR) to enter the folate cycle^10,11^. Both SFA and 5-CH_3_-THF are naturally occurring in humans, but 5-CH_3_-THF is the predominate form of folate in cerebral spinal fluid, blood serum, and erythrocytes^11,12^. Pre-conceptional FA supplementation to women of childbearing age can decrease the occurrence of NTDs by up to 85 percent^13,14^. The daily recommended dose of 400μg is reported to provide sufficient FA to produce optimal serum and red blood cell (RBC) folate concentrations in pregnant women^15,16^. Unfortunately, FA supplementation started after the first trimester of pregnancy, does not prevent NTDs. In response to this concern, FA fortification of grains and cereal products has been mandatory in the United States since 1999, and post-fortification monitoring indicates that this has been effective in decreasing the occurrence of NTDs^17-19^.

## Folate Receptors

Transport of folates into cells is principally accomplished via folate receptors (FR). There are four human folate receptor isoforms (FOLR1/FR-α, FOLR2/FR-β, FOLR3/FR-γ, FOLR4/FR-δ). Initially isolated from human placenta, the folate receptor isoforms have been shown to have tissue-specific and cell-specific expression patterns^20-23^. The folate receptor isoforms are externally bound to the plasma membrane by glycosyl-phosphatidylinositol (GPI) anchors, but a high-affinity folate receptor has also been observed unbound without GPI modifications in plasma. Folate binding data and immunodetection of FOLR1/FR-α indicates expression is primarily found in the plasma, placenta, choroid plexus, and the brush border membranes of the kidney^24,25^. In the developing mouse embryo, Folr1 is essential for embryonic and neural tube development^26^, and *Folr1* is observed to be highly expressed in yolk sac, neural folds, and the neural tube^27^. The Folr2 protein is non-essential for development in mice, as Folr2 knockout mice develop normally^26^, and human FOLR3 lacks a mouse homolog. FOLR4/FR-δ, also known as Juno, is a member of the folate receptor family but is not involved in folate uptake^28^. JUNO is located on the surface of mammalian eggs and recognizes a sperm-counterpart, IZUMO1, to facilitate fertilization^29^.

## FOLR1 Antagonism and Birth Defect Risks

The developmental toxicity of anti-folates was first described in chickens^30^. Subsequent studies in mice, rats, rabbits, and monkeys demonstrated embryotoxicity and teratogenicity that varied based on developmental timing and dose^31-33^. Anti-folate, e.g. methotrexate (MTX), developmental toxicity and the mitigation or rescue by supplemental folate has been demonstrated in experimental animals and in clinical practice, respectively^34,35^. As immunotherapy began to also target FOLR1-positve tumors, antibodies against FOLR1/ FRα were reported to antagonize or block cellular folate uptake into placenta cells^36,37^. The initial report of FOLR1 autoantibodies examined FOLR1 antiserum on embryogenesis in a rat model. It was demonstrated that antiserum against FOLR1/FRα and FOLR2/FRβ caused dose-dependent developmental toxicity^36^. Large doses of antiserum caused 100% embryo-lethality, while lower doses increased the incidence of NTDs in exposed offspring. Autoantibody directed against human FOLR1/ FRα were also detected in human serum in association with an increased risk of NTDs^37^. We developed a sensitive assay to identify FOLR1-folate binding interactions with autoantibodies^38^. Since the initial report, we have examined several population cohorts and have found that autoantibodies to FOLR1 are significantly higher in NTD-affected pregnancies compared to control subjects, supporting the association between FOLR1 autoantibodies in maternal serum and increased risk of folate-responsive birth defects, including NTDs, in expectant mothers^39-41^.

## DTG and FOLR1 Antagonism

Based on the clinical benefit of FA in the prevention of birth defects in humans, the utility of folates in rescuing embryo lethality and NTDs in FOLR1 mutant mice^26,42^, and the risk of NTDs with DTG exposure or FOLR1 antagonism, we hypothesized that DTG is a FOLR1 antagonist. This report represents the first published data on the testing of antagonism of FOLR1-folate binding by DTG. It utilizes a vertebrate animal model (*Danio rerio*) to evaluate folic acid as a major modifier of DTG-induced developmental toxicity. Developmental toxicity testing of antifolates directly on pregnant women is medically unethical, so an *in vitro* cellular model (HTR-8/SVneo) was used to study human placental-trophoblast interactions with folate in the presence of DTG. Although various cell types are recognized in embryonic placenta primordia and term placentas, trophoblasts were selected for testing, because they perform several essential functions for viviparous development, have essential roles in folate uptake, protect the embryo/fetus from the immune system and chemical insults, and provide bidirectional nutrient/waste flow required for embryonic and fetal growth, development, and maturation^43-45^. These experimental models allowed for the rapid determination of DTG-induced developmental toxicity and are of critical importance to the HIV, ART, and birth defect research communities. Complex time-dependent protein-ligand-drug interactions modify embryo susceptibility to the developmental toxicity of DTG. The identification of these biochemical interactions and resultant developmental toxicity has clinical and biological significance beyond this report, as the results from these studies are expected to inform and guide future animal models and clinical studies of DTG-based ART in women of childbearing age.

## RESULTS

Standard FOLR1-folate curves were produced in the absence of DTG (Figure 1). In order to test for interactions between DTG and FOLR1, curves were produced in the presence of varying concentrations of DTG (prepared as 1:2 dilutions from 100μM-0.73125μM). Results indicated that FOLR1-FA-HRP signal decreased in the presence of clinically relevant concentrations (3-10 μM) of DTG, and the response was concentration dependent (Figure 1). Resultant intensity data for DTG-FOLR1-FA-HRP were normalized to comparable antagonism by FA. The resulting curves (Figure 1B, 1C) indicate that DTG is a partial antagonist of FOLR1-folate binding. The reported therapeutic range of DTG serum concentrations are 3-10μM^46,47^, and DTG produces partial inhibition within this range, with an unadjusted IC_50_ of approximately 4.4μM (Figure 1a, non-linear best fit).

**Figure 1.**
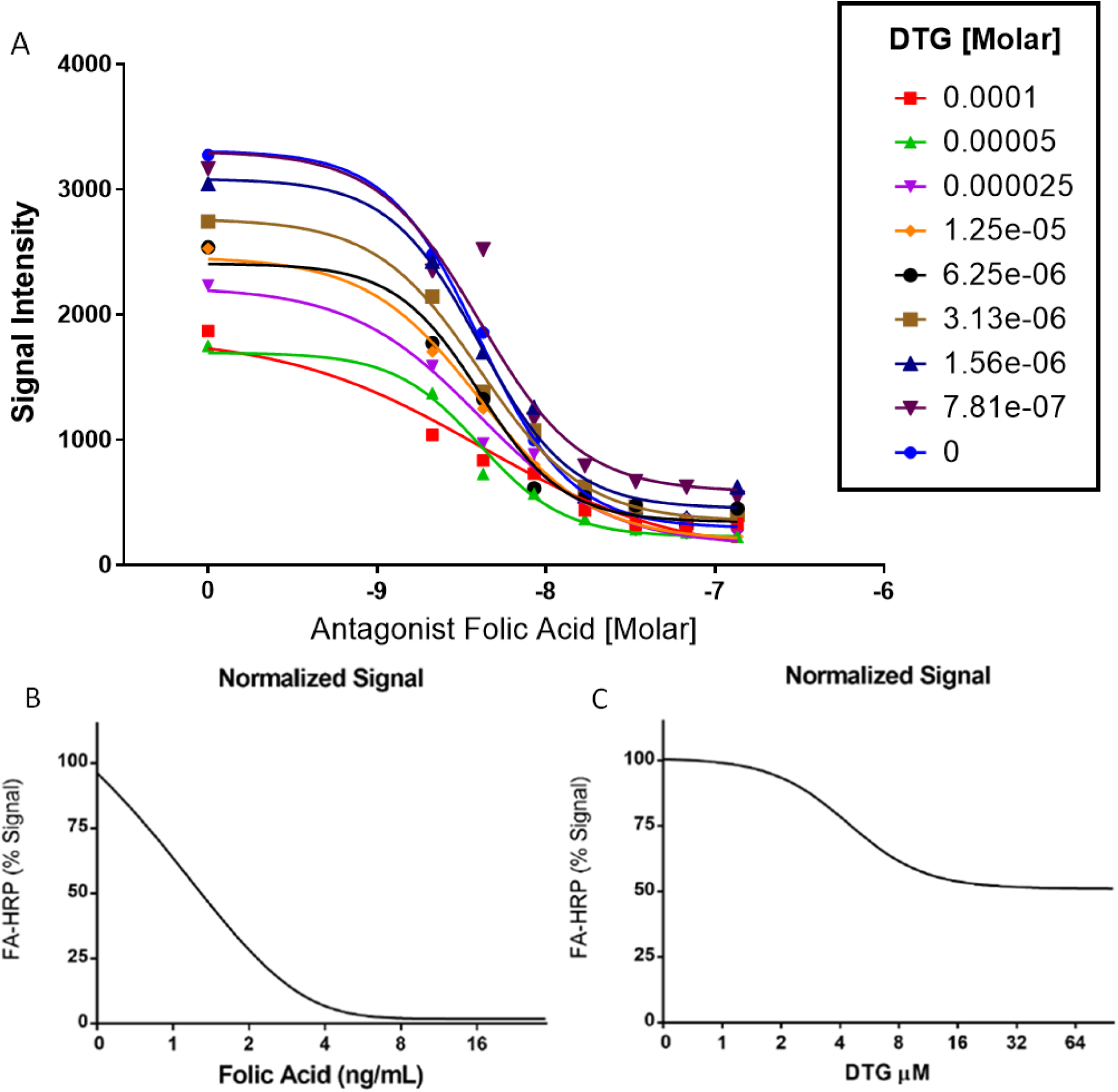
Binding Curves for DTG Antagonism of FOLR1. A) The binding curves produced by DTG and FOLR1-folate show concentration-dependent decreased signal intensities. Maximum FA-HRP is decreased and curves are marginally shifted left in a concentration-dependent manner (P=0.06). A left shift is expected with competitive antagonists. These data are most consistent with partial or non-competitive antagonism. B) Normalized analysis of FA (competitive) binding to FOLR1. C) Normalized analysis of DTG indicates partial antagonist activity (56% signal at 16μM) by DTG, showing allosteric noncompetitive antagonism of FOLR1 by DTG.

We determined calcium (2mM) increased folate-FOLR1 binding in cells and in protein binding assays. Initially, we examined cations in cell culture media (Ca^+^2, Fe^+^3, Mg^+^2, and K^+^), yet absent in DBPS, for modification of DTG-FOLR1-folate interactions. These FA-HRP binding studies (Supplemental 1) with DPBS, 1% FBS, and 20μM DTG on HELA cells indicated cations increased folate binding 10.21% (P = 0.0655) for K^+^, 10.75% (P = 0.0456) for Mg^+^2, 10.84% for Fe^+^3. 10.75% (P = 0.0480), and 59.96% (P < 0.0001) for Ca^+^2. Analysis by ANOVA (P < 0.0001), with post-hoc Dunnett’s. The modification of folate binding with calcium alone or in the presence of DTG was unexpected, so we tested calcium (2mM) in the microtiter assay. Calcium increased folate-FOLR1 interactions (+42.3%) in the absence of DTG (Figure 2) compared to control (no DTG, no calcium). Low concentrations of DTG (0.25-4μM) in the presence of calcium (2mM) resulted in additional increased folate binding (+116%, 4μM), but as the concentrations of DTG increased above 4 μM, 4-32μM, there was a decreased folate binding (32μM, - 23%). Similarly, there was increased folate binding to cells in the presence of DTG (20μM) when Ca^+^2 (2mM) was present (Figure 3). This response occurred with nuclei contraction and decreased cell number, which indicates cytotoxicity and cell-loss (Figure 3).

**Figure 2.**
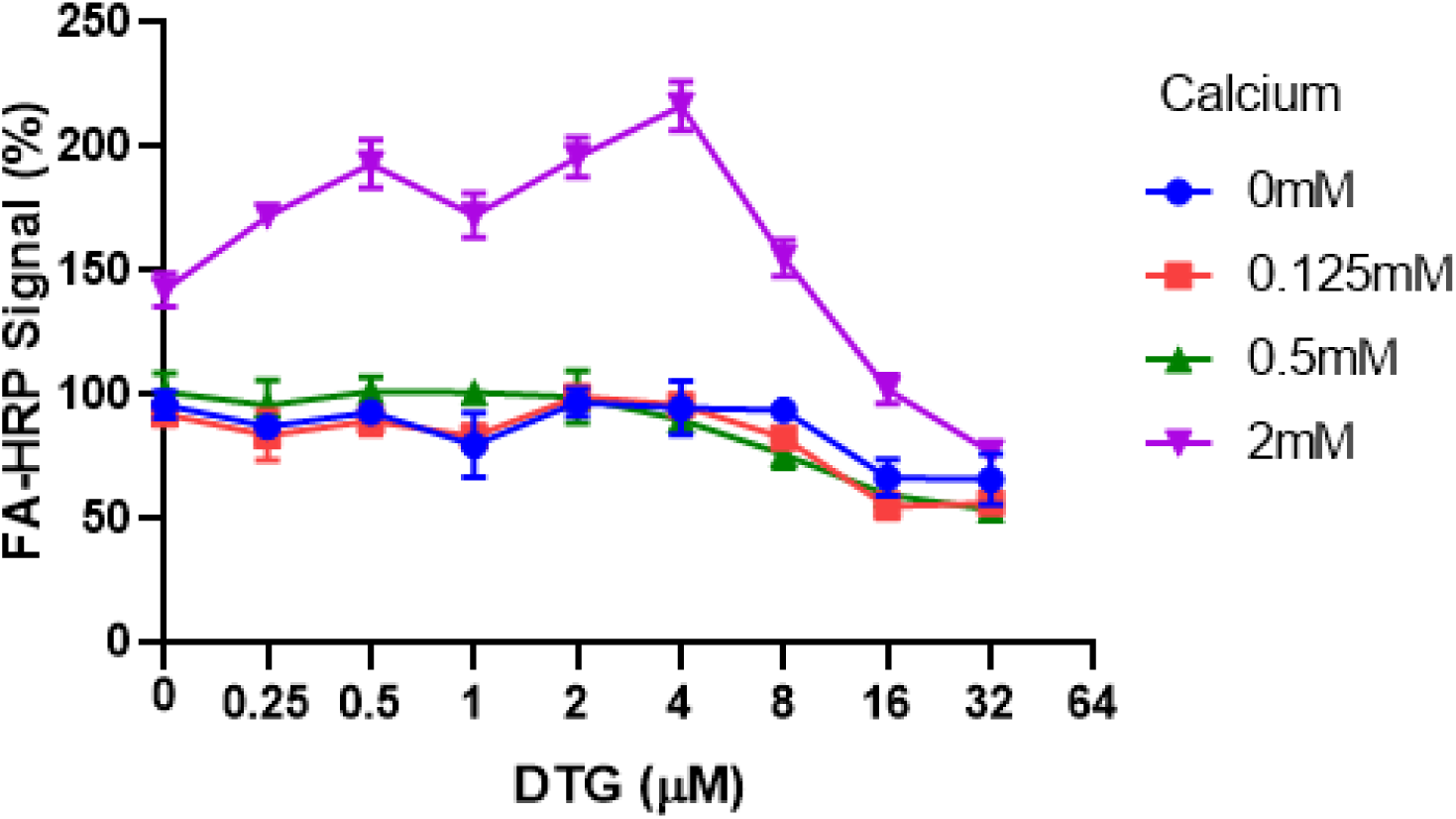
Calcium Modifies Folate Binding to FOLR1 and DTG Interactions. Calcium (2mM) modified folate binding to FOLR1 in the microtiter protein-ligand binding assay. In the presence of Ca^+^2 (2mM) and absence of DTG, folate receptor bound 42.3% more folate (FA-HRP). The response to DTG in the presence of 2mM Ca^+^2 appeared bi-phasic. The DTG-Ca-FOLR1 interaction below 4μM increased folate binding, but at concentration above 4μM, the binding fell to 77%, or decreased −23% compared to untreated cells.

**Figure 3.**
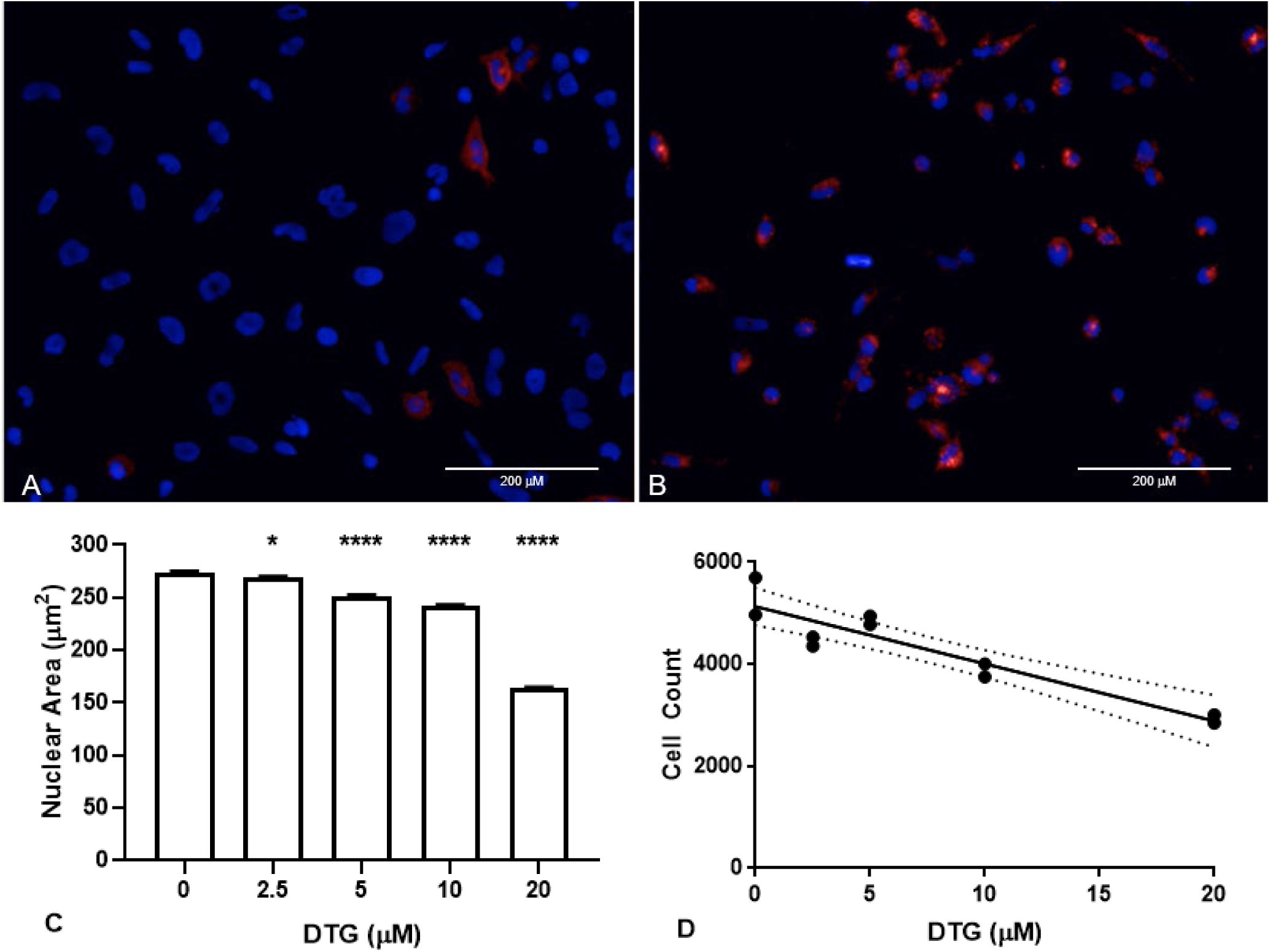
Calcium Alters DTG-FA-680 Interactions in HTR-8/SVneo. Images of HTR-8/SVneo cells exposed to 20μM DTG, 50nM FA-680 (red), and Hoechst 3342 (blue, live-nuclei) in buffer (1X DPBS, 1% FBS) (A) without additional Ca^+^2 or (B) with 2mM Ca^+^2 for 1 hour. Bar = 200μm. After 24hrs in RPMI (1% FBS, 50nM FA-680) and variable concentrations of DTG, nuclei showed DTG dependent (C) nuclear contraction (* P = 0.0285, **** P < 0.0001) and (D) decreased cell counts (r = −0.9318, R^2^= 0.8683, P < 0.0001, dotted lines show 95% confidence interval).

Additional competitive binding assays using relevant folates (FA, 5-CH_3_-THF, SFA) and folate receptors (FOLR1, FOLR2, bovine folate binding protein (bFBP) were performed to compare competitive binding. Results indicate that 1.5 ng/mL of folic acid is the equivalent inhibition produced by approximately 50ng/mL of 5-CH_3_-THF with FRα /FOLR1 (Supplemental Table 1). It is important to note that the predominate folate in biological fluids and tissues is 5-CH_3_ THF (5-15 ng/mL), and the comparable IC_50_ of 1.5ng/mL FA is equivalent to approximately 50ng/mL 5-CH_3_-THF.

The DHFR enzyme activity assay was performed, and the resulting absorbance (340nm) reading produced linear slopes of enzyme kinetics over time (Figure 4). Inhibition of DHFR by MTX (positive control) from 100-10μM produced slopes that ranged from −3.42 to −5.12, and in the absence of inhibition (negative control) there were steep linear kinetics (−21.36 to −28.17). There was no observed change in DHFR enzyme kinetics (range of −20.59 to −28.16) with DTG (25-100μM). These data do not support inhibition of DHFR by DTG; so DTG does not inhibit downstream metabolism by DHFR, as observed with MTX.

**Figure 4.**
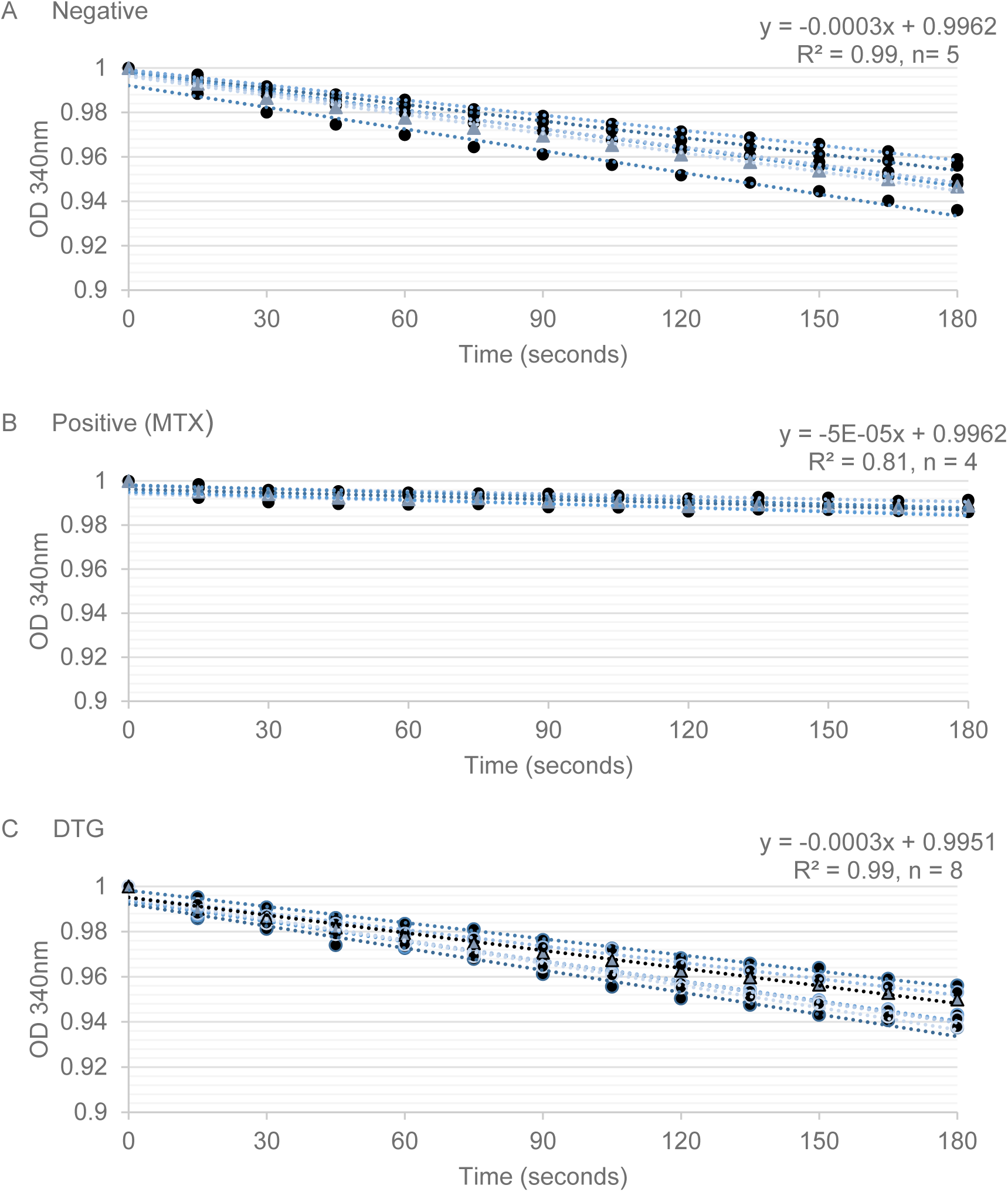
DHFR Activity in the Presence of MTX or DTG. DHFR catalyzed NADPH-dependent reduction of DHF to THF, reducing 340nM optical density (Y-axis, OD) readings over time (X-axis, seconds) to produce enzyme kinetics (linear slope). The initial OD reading was set to 1 OD, and the average OD change for each treatment over time was used to calculate the linear-fit equation. A) In the absence of an inhibitor, the reaction proceeded with a slope of −0.003 (OD/seconds). B) MTX (100μM) showed inhibition of DHFR activity, as observed by a slope approaching zero (−5e-05). All DTG concentrations (100-12.5μM) tested resulted in an enzymatic slope comparable to those observed in the negative control (−0.003) and are consistent with no DHFR inhibition.

The results obtained in the zebrafish model demonstrate the presence of developmental toxicity due to early embryonic exposure to DTG (Figure 5). Toxicity was observed following DTG exposures beginning at 1-, 3-or 5-hours post-fertilization (hpf) (Figure 5, Figure 6, and Supplemental Table 2). In contrast, limited developmental toxicity was observed when exposure began during gastrulation (6-8 hpf) (Figure 6). The specificity of DTG developmental toxicity was tested via rescue of DTG-induced developmental toxicity by supplemental folate (Figure 6, Supplemental Table 2). When DTG exposure started at 1 or 3 hpf, high developmental toxicity (80-100%) was observed at 24 hpf. Developmental toxicity was rescued by co-exposure to folate (0-7.69% mortality). When DTG exposure started at 5 or 6-8 hpf, developmental toxicity decreased as hpf increased, with the lowest toxicity at 6-8 hpf (Figure 6).

**Figure 5.**
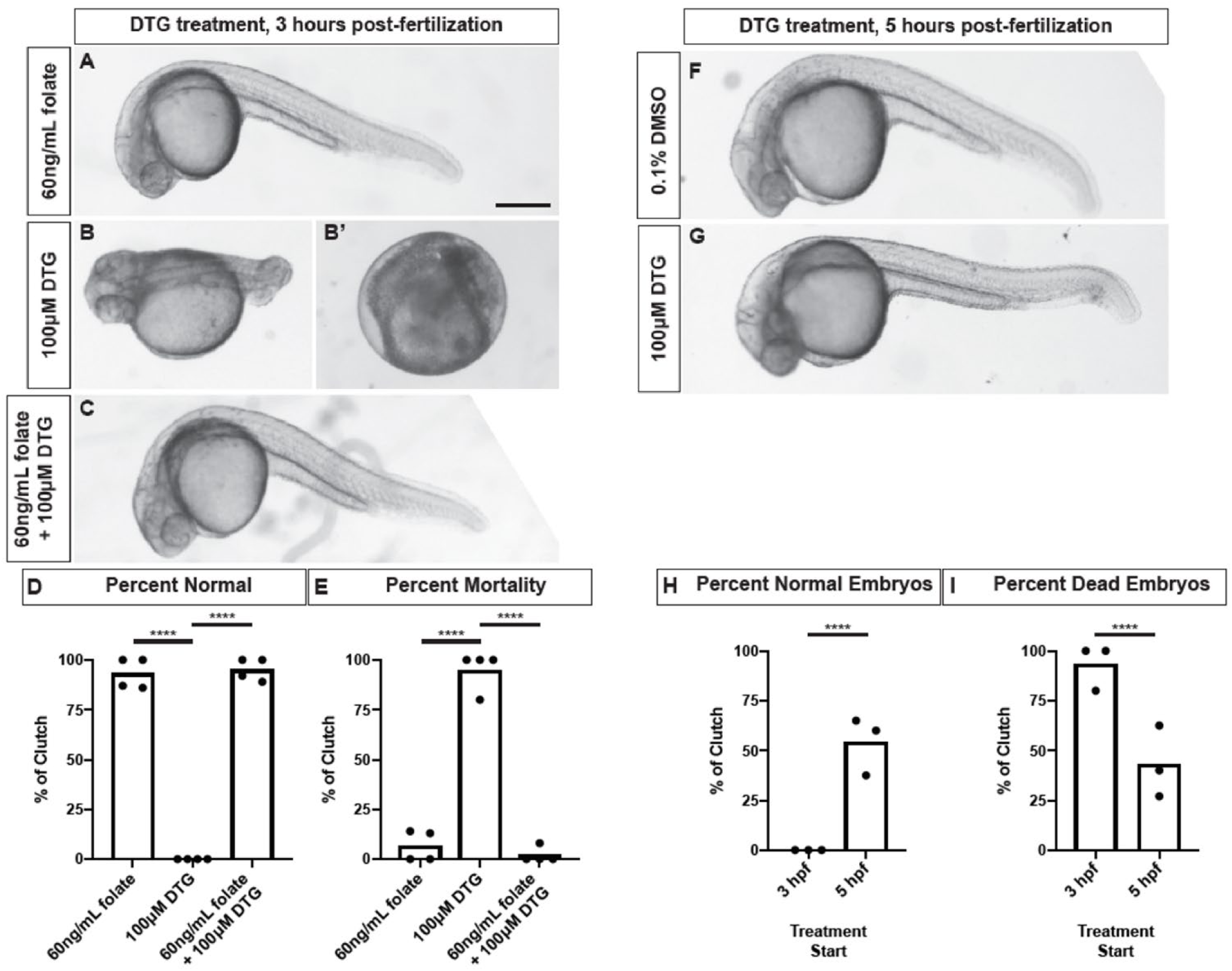
Folic Acid Rescue of DTG Developmental Toxicity. Folate exposure rescues DTG toxicity in zebrafish embryos. (A-C) Representative images of zebrafish embryos exposed to folate and/or dolutegravir (DTG) starting at 3 hours post fertilization (hpf). Embryos were imaged at 1-day post fertilization. B-B’ represents DTG-induced toxicity in embryos that is rescued upon co-treatment with folate (C). (D-E) Quantification of normal (D) and dead (E) embryos following folate, DTG, or DTG+folate treatment starting at 3 hpf. Each dot represents the mean percent from a clutch of embryos. Bars represent the average of 4 clutches. DTG exposure induces 100% toxicity in embryos while co-exposure with folate rescues toxicity. (F-G) Representative images of zebrafish embryos exposed to vehicle (DMSO) or DTG starting at 5 hpf. Embryos were imaged at 1-day post fertilization. (H-I) Quantification of normal (H) and dead (I) embryos following vehicle or DTG treatment starting at 3 or 5 hpf. Each dot represents the mean percent from a single clutch of embryos. Bars represent the average of 3-4 clutches. The 3 hpf clutches are the same groups from panels B, B’, D, E. DTG exposure is significantly reduced when starting exposure at 5 hpf compared to 3 hpf.

**Figure 6.**
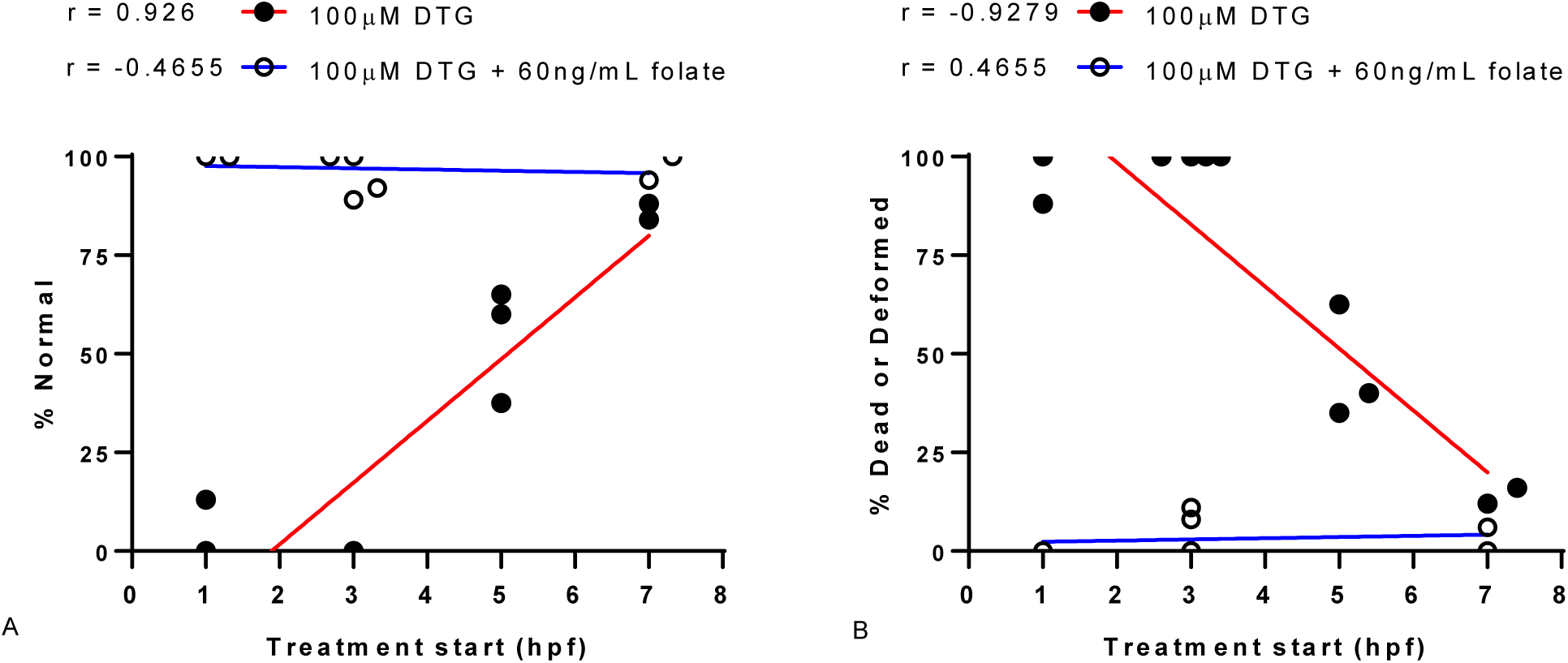
Critical Period of DTG Developmental Toxicity in Zebrafish. The critical window of exposure for DTG was determined over time by testing the start of DTG exposure at different times (1, 3, 5, and 6-8 hpf). For analysis, the 6-8 hpf range was plot at 7hpf. The normal phenotype (% normal, A) and mortality-morbidity (% dead or deformed, B) observed at 24 hpf was plot as a function of DTG treatment (100μM) start time. Linear regression was fit to the resultant data (R^2^= 0.86, P < 0.05, for DTG A & B). The mortality-morbidity for DTG exposure was highest with early pre-gastrulation exposures (1-3hpf), differed significantly over time (1-3 hpf vs 5-8hpf, T-test P = 0.001), and was lowest at 6-8hpf. Co-exposure of folic acid (FA, 60ng/mL) with DTG (100μM) results in rescue from developmental toxicity.

## DISCUSSION

Folic acid fortification and supplementation are the major nutritional modifiers for NTD risk reduction. The function of FOLR1 is essential to embryonic development, neurodevelopment, and neural tube closure. Folate antagonists, autoimmunity to FOLR1, or deletion of *Folr1* in mice increases the risk of developmental toxicity, birth defects, and NTDs. Dolutegravir is a partial antagonist of FOLR1 (Figure 1), and early embryonic exposure to DTG produces developmental toxicity in zebrafish that can be rescued by supplemental FA (Figure 5, Supplemental Table 2). Folic acid is a ligand and competitive antagonist of FOLR1-FA-HRP. Dolutegravir shows partial antagonist activity, lacking the ability to saturate binding, with no concentration dependent signal changes above 16 μM (Figure 1B, 1C). A concentration of 10-12μM DTG produces normalized inhibition comparable to the IC_50_ of 1.5 ng/mL for FA. The normal serum concentration range of folates is 5-15 ng/mL, so a comparable IC_50_ of 1.5ng/mL folic acid appeared small; however, FA is a higher affinity ligand for folate receptors than other folates. Whether other folates (e.g. SFA or 5-CH_3_-THF) can rescue DTG developmental toxicity and with potentially greater efficacy is unknown, but it is expected that the IC_50_ results (Supplemental Table 1) based on relative competitive binding to FOLR1 will correlate with concentrations needed to rescue DTG developmental toxicity. Based on binding data, consistent findings of increased mortality in zebrafish embryos exposed to DTG, and rescue of DTG developmental toxicity by FA, we conclude that DTG is teratogenic due to FOLR1 antagonism.

We utilized human placental trophoblast cells (HTR-8/SVneo) exposed to DTG at clinically relevant concentrations as an *in vitro* model of first trimester placental cell interactions with DTG and folate. Adverse interactions with DTG have been reported clinically with co-administered polyvalent cations, including calcium and iron (Tivicay, 09/2018, Summary of Effect of Coadministered Drugs on the Pharmacokinetics of Dolutegravir). The clinical concern is that co-administration can result in decreased DTG uptake and lower serum concentrations^48^. The increase of folate binding in cells was unexpected based on the FOLR1-folate binding microtiter assay. The increase in folate-FOLR1 interactions (+42.3%) produced by calcium alone appears biologically relevant to folate-FOLR1 interactions. This interaction is supported by another report that indicated calcium chloride (1mM) resulted in doubling of folate binding to rat intestinal mucosal cells^49^. Based on a red blood cell study that reported DTG produces cytotoxicity in the presence of extracellular calcium^50^, we focused on calcium as a modifier of folate binding in HTR-8 cells and in the FOLR1 microtiter binding assay. The microtiter assay reproduced the increased folate binding observed in the cell culture assay if calcium (2mM) was included in the binding assay. These data indicate that calcium is a modifier of folate binding to folate receptors, and we propose DTG chelation of calcium modifies FOLR1 interactions and cytotoxicity (Figure 3).

Mechanistically, DTG binds to magnesium at the active site of the HIV integrase enzyme, thereby preventing integration of viral DNA into the host genome. Drugs in this class are unfortunately susceptible to cation interference if co-administered with mineral supplements or antacids^48^. These results indicate the increased folate detected in the trophoblast model is a result of DTG-Ca-FOLR1-folate interactions, and the increase in signal represents localized increased concentrations of DTG and folate due to interaction with FOLR1. As soluble calcium increases folate-FOLR1 interactions and also interacts with DTG, the resulting chelation and FOLR1 interaction decreases DTG-Ca-FOLR1-folate solubility. This model is consistent with nuclei contraction and deceased cell numbers observed when DTG was coadministered with calcium and folate (Figure 3). Based on these data, we propose the increase in FOLR1-folate binding in cells exposed to DTG is due to DTG binding of calcium, and calcium-responsive FOLR1-folate binding, producing DTG-FOLR1-folate or DTG-Ca-FOLR1-folate complexes with limited DTG-folate solubility and increased cytotoxicity. The FOLR1 epitopes involved in DTG or Ca^+^2 binding, structural and stochiometric interaction of these molecules, and testing of other integrase inhibitors for similar interactions are open topics for future studies.

At 24 hpf, most DTG exposed embryos were dead clumps of cells, so future studies may also consider looking at DTG-induced zebrafish phenotypes in greater detail to precisely determine which cell populations are affected and when pathologies or malformations occur. We observed that when exposure began during gastrulation (6-8hrs post-fertilization), there was decreased developmental toxicity (Figure 6). This result is consistent with and supports the principle of a critical period for developmental toxicity (1-4 hours, or < 6-8 hours post fertilization). Wilson indicated, “susceptibility to teratogenesis varies with the developmental stage at the time of exposure to an adverse influence”^51^. The critical period of exposure/toxicity to early embryos is also consistent with clinical reports on DTG with increased risk for NTDs ^2^. Specifically, late gestational exposures to DTG have not been associated with adverse developmental outcomes. Developmental timing and dosages, for FOLR1-antagonists and folates, should all be considered major modifiers of developmental toxicity risk in animal models and humans. These data may also partial explain the lack of statistically significant preclinical teratogenicity. Specifically, in rats and rabbits, DTG was administered during organogenesis, days 6-17 and 6-18, respectively. This is consistent with Guidelines for Industry (FDA) and guidelines put forth by the International Conference on Harmonization, but a critical period from 1-4 hours in zebrafish parallels the blastula, pre-gastrulation, pre-implantation and implantation periods of mammalian embryonic development (< 3-weeks in humans, <6.0 days post conception in mice and rats).

The mechanisms by which NTDs and congenital malformations are produced are varied, but folate antagonism is an accepted teratogenic mechanism of action in animal models and humans^52,53^. During our review of preclinical studies, we found there was an NTD in the preclinical study at 40 mg per Kg DTG in rabbits (Oral Study for Effects of S-349572 Sodium on Embryofetal Development in Rabbits). While one affected fetus or litter in a teratology study is not statistically significant, it is worth mentioning due to recent human studies. We propose the combination of fortified folate diets coupled to a critical period in zebrafish that parallels pre-implantation, implantation, and early gastrulation stages in mammalian embryos (Carnegie Stage < 8) limited previous detection of developmental toxicity.

We present the specificity of DTG developmental toxicity by rescue with folic acid, which is known to be effective in folate responsive anemia, increasing average serum folate, and decreasing NTD risk. The FA rescue of DTG developmental toxicity was proposed because FA has the highest affinity to FRs (Supplemental Table 1) and is commonly used in food fortification strategies. Clinically, SFA is preferred to decrease the anti-folate toxicity of MTX, because the use of SFA bypasses DHFR activity, which is inhibited by antifolates/FOLR1 antagonists. However, we determined that unlike known antifolates, DTG does not inhibit DHFR (Figure 4). Collectively, these data are consistent with partial antagonist activity at FOLR1, such that the binding of FOLR1 is altered but still interacts with FA in the presence of DTG, but the folate binding and enzymatic activity of DHFR is unchanged by DTG. Accordingly, DTG is not proposed to directly interact with folate or the binding domain of FOLR1 or DHFR, but additional structural analysis will be needed to identify and provide the location for the antagonist-protein interaction between DTG and FOLR1.

We have previously reported that homozygous deletion of *Folr1* in mice is lethal, but folate supplementation of *Folr1* mutants produces a range of developmental defects that vary by genetic background and range from lethality, to NTDs, to complete rescue^26,42^. Based on these results, we propose the recommended FA (2-3 mg/kg) content in laboratory animal chow masked the developmental toxicity of DTG. We propose future mammalian animal testing of DTG developmental toxicity is warranted, but pre-, peri-, and post-implantation DTG exposure should be explored in studies using both low-folate diet (e.g. 0.3 mg/kg) and standard-folate diet to examine folate responsive defects and folate masking of DTG developmental toxicity. Furthermore, as more human cohort data is generated, or additional clinical studies are performed on DTG, biological interactions and clinical correlations between natural folate intake, folic acid fortified diets, supplemental folate, calcium intake, and resultant DTG, folate, and calcium in blood and serum and should be investigated. It has been previously reported that in human populations where folate fortification is in place, the prevalence of folate deficiency is approximately 0.1% or 1 per 1,000 individuals, but in unfortified population, the base line incidence may be > 20%, 1 in 5, or possibly higher^54^. This 100 to 200-fold difference in the incidence of folate deficiency lowers the average serum folate, increases NTD risk, and may also increase the risk of NTDs associated with early gestational *in utero* DTG exposure. According to a WHO review from 1993 to 2005, 42% of pregnant women had iron-folate related anemias worldwide and almost 90% of anemic women reside in Africa or Asia^55^. On the other hand, preclinical animal studies are expected to approach 0% folate-deficient on standard diets, because they all have adequate folate intake. The fortification of enriched cereal-grain products (mandatory in the U.S. on January 1, 1998) was intended to increase folate intake among childbearing-aged women to reduce their risk of NTD-affected pregnancies, but vitamins and minerals can also mitigate other environmental and pharmaceutical toxicities. The data reported in this study indicate that the research and medical communities should consider folic acid supplementation and maternal folate status as a major modifier of DTG-induced developmental toxicity.

## METHODS

### Folate Binding Assays

A microtiter binding assay was used to measure inhibition of FOLR1-folate binding by DTG, as previously described for FOLR1 autoantibody^38,41^. Briefly, FOLR1 (1 μL at 50ng/μL) was immobilized on microtiter plates in bicarbonate buffer (50mM, pH 9.0). The following day, unbound protein is removed by washing thrice with tris-buffered saline (pH 7.6). Ligand in the form of FA is used to compete with labeled (e.g. horse radish peroxidase) folic acid (FA-HRP) for competitive binding to FOLR1. Standard dilution (1:2) of FA (60.0 to 0.8125ng/mL, 0ng/mL) results in FOLR1-folate binding curves. Additional competitive binding assays using relevant folates (FA, 5-CH_3_-THF, and SFA) and folate receptors (FOLR1, FOLR2, bovine folate binding protein (bFBP)) were performed to compare binding of different folates to different folate receptors.

Reference pharmacokinetic data of DTG in humans indicates a single daily 50 mg dose of DTG produces blood concentrations ranging (Cmin-Cmax) from 3-10 micromolar ^46,47^. In order to test for interactions between DTG and FOLR1-folate, standard FOLR1-folate curves were produced, as described above, in the presence of varying clinically relevant concentrations of DTG (prepared as 1:2 dilutions from 100 μM-0.73125 μM). Base on the results of *in vitro* studies, calcium was also examined for interactions with FOLR1-folate binding by DTG. Calcium was tested by adding calcium chloride (0.5-2 mM) to the competitive binding buffer (DTG, FA-HRP).

### In vitro Testing of DTG Development Toxicity

The impact of DTG on folate binding was measured using a transformed first trimester human trophoblast cell line (HTR-8/SVneo) (PMID: 7684692), provided by Jacqueline Parchem, M.D. (Department of Obstetrics and Gynecology, Baylor College of Medicine). The HTR-8/SVneo cell line (ATCC, CRL-3271) was cultured under standard culture conditions (37°C, 5% CO_2_) using RPMI-1640 with 5% fetal bovine serum (FBS). The evening before testing, cells were plated (5,000 cells per well) into 96-well plates (Corning 3630). Live cell imaging using near-infrared-labeled folate (FolateRSense 680, Perkin Elmer), prepared in Dulbecco’s phosphate-buffered saline (DPBS) with 1% FBS, was used to determine the impact of DTG on folate interactions with human trophoblast cells.

We examined Ca^+^2, Fe^+^3, Mg^+^2, and K^+^, prepared from calcium chloride, iron III sulfate, magnesium sulfate, or potassium chloride for modifications of DTG-FOLR1-folate interactions. The concentrations of Ca^+^2 (1.8mM), Fe^+^3 (250nM), Mg^+^2 (810μM), and K^+^(5.3mM) tested are typical for cell culture media or serum. For example, normal extracellular concentrations of calcium are 1-3 mM and intracellular concentrations are 0.1-0.2 µM, so calcium chloride (0-2mM) was added to DPBS with 1% FBS. Folic acid (0-60ng/mL) and DTG (0-100μM) were likewise dissolved in DPBS with 1% FBS for testing. Following incubation (1 hour, PBS 1% serum) with test compounds, cells were washed in DPBS with 1% FBS. Live-cell nuclei were stained using Hoechst 33342 (1 μg/mL, 10 minutes) (Thermo Fisher Scientific). Images were captured and analyzed on the Operetta High Content Imaging System (Perkin Elmer). Folate uptake was also determined by using unlabeled folic acid, without FolateRSense 680, in the presence of test compounds. These cells were washed in DPBS with 1% FBS and unlabeled folate concentrations determined in cell lysate (1X Tris buffered saline, ascorbic acid, 0.1% Tween20) using the microtiter folate binding assay.

### DHFR Activity

Dihydrofolate reductase catalyzes the reduction of FA into dihydrofolate (DHF), and DHF into tetrahydrofolate (THF). Blocking DHFR by anti-folates (e.g. MTX) can induce cell death via inhibition of nucleoside synthesis and thereby nucleotide and DNA/RNA synthesis^56^. Based on the observed anti-folate activity of DTG on FOLR1, we hypothesized that DTG would inhibit downstream DHFR activity. DHFR activity and screening of DHFR inhibitors testing was examined using the Dihydrofolate Reductase Assay Kit (Sigma). The assay is based on the ability of DHFR to catalyze the reversible nicotinamide adenine dinucleotide phosphate (NADPH) dependent reduction of DHF to THF. The reaction progress is followed by monitoring the decrease in concentration of NADPH by absorbance at 340 nm in the presence of enzymes and substrates: DHF + NADPH + H+ ↔ THF + NADP+. The kit included MTX as a positive control, and the assays were performed as described by the manufacturer.

### In vivo Testing of DTG Development Toxicity in Zebrafish

Adult zebrafish were raised at 28.5°C on a 14-h light, 10-h dark cycle using an Aquaneering recirculating water system (Aquaneering, Inc., San Diego, CA). All zebrafish used for experiments were wild-type AB strain^57^. All procedures were approved by the BCM Institutional Animal Care and Use Committee. Adult zebrafish were allowed to spawn naturally in groups. Embryos were collected in intervals of 10 minutes to ensure precise developmental timing and staged following collection, placed in 60 cm^2^Petri dishes at a density of no more than 100 per dish in E3B media (60X E3B: 17.2g NaCl, 0.76g KCl, 2.9g CaCl_2_-2H_2_O, 2.39g MgSO_4_ dissolved in 1L Milli-Q water; diluted to 1X in 9L Milli-Q water plus 100 μL 0.02% methylene blue), and then stored in an incubator at 28.5°C on a 14-h light, 10-h dark cycle until treatment.

Folate receptor expression and anti-folate toxicity has previously been examined in zebrafish embryos^58,59^. Embryonic responses to DTG and folate were tested based on previously published MTX developmental toxicity testing in zebrafish^58^. Fertilized embryos were placed in 6-or 12-well plates (15-21 embryos/well) in 2-3 ml of embryo medium per well, followed by chemical treatment beginning at 1, 3, 5, and 6-8 hours post fertilization (hpf) with daily renewal. The embryos were incubated at 28.5°C with a 14/10 hr light/dark cycle and assessed for developmental malformations and lethality at 24 hpf by light microscopy. For zebrafish treatments, stock solutions were prepared at 100mM, in DMSO. Working solutions were diluted 1000X into E3B resulting in a final concentration of 0.1% DMSO for static exposure. DTG was tested at 100 μM for preliminary experiments, which is approximately 10X C_max_ from clinical testing. Folic acid supplementation was 60ng/mL, prepared in water at 60μg/mL and diluted to 60ng/mL in E3B for testing. Zebrafish embryo phenotypes were visually analyzed using light microscopy and scored as an average percentage of embryos with mortality or malformation out of all treated embryos (5-26 embryos/ treatment) from 2-4 experimental replicates.

## SUPPLEMENTAL TABLES

**Supplemental Table 1.** Folate Receptor IC_50_ Interactions with Folate Ligands Binding curves were prepared for folates and isolated folate receptors. Soluble bovine folate binding protein (bFBP) from milk shows the highest affinity to all ligands. The inhibition of 1.5ng/mL FA by DTG is comparable to inhibition of 50 and 90ng/mL for 5-CH_3_ THF for FOLR1 and FOLR2, respectively.

| Ligand | IC 50 (ng/mL) |  |  |
| --- | --- | --- | --- |
|  | FOLR1 | FOLR2 | bFBP |
| Folic Acid (FA) | 1.45 | 1.46 | 0.26 |
| 5-CH<sub>3</sub> THF | 50 | 90 | 10 |
| S-Folnic Acid (SFA) | 20900 | 56110 | 270 |

**Supplemental Table 2.** Folic Acid Rescue of DTG Developmental Toxicity Folate exposure rescues DTG toxicity in zebrafish embryos. A) Embryos were statically exposed beginning at 3 post fertilization (hpf) to dolutegravir (DTG), folic acid (FA), or DTG + FA. High developmental toxicity (80-100%) were observed with DTG treatment, with rescue by co-exposure to folate (0% mortality). B) Embryos were statically exposed beginning at 3-hour post fertilization to DTG, FA or DTG + FA. Tables report number of dead and normal embryos for clutches; embryos in different clutches have different parents and are therefore biological replicates.

| Treatment | Clutch | 24 hpf |  |  |  |
| --- | --- | --- | --- | --- | --- |
|  |  | Dead | Moribund | Normal | Mortality |
| 100uM DTG | 1 | 9 | 0 | 0 | 100.00% |
|  | 2 | 8 | 2 | 0 | 80.00% |
|  | 3 | 15 | 0 | 0 | 100.00% |
|  | 4 | 12 | 0 | 0 | 100.00% |
| 60ng/mL FA | 1 | 1 | 0 | 4 | 20.00% |
|  | 2 | 1 | 0 | 6 | 14.29% |
|  | 3 | 0 | 0 | 5 | 0.00% |
|  | 4 | 2 | 0 | 13 | 13.33% |
|  | 5 | 0 | 0 | 12 | 0.00% |
| 100uM DTG +<br>60 ng/mL FA | 1 | 0 | 1 | 8 | 0.00% |
|  | 2 | 0 | 0 | 10 | 0.00% |
|  | 3 | 0 | 0 | 16 | 0.00% |
|  | 4 | 1 | 0 | 12 | 7.69% |

## SUPPLEMENTAL

**Supplemental Figure 1.**
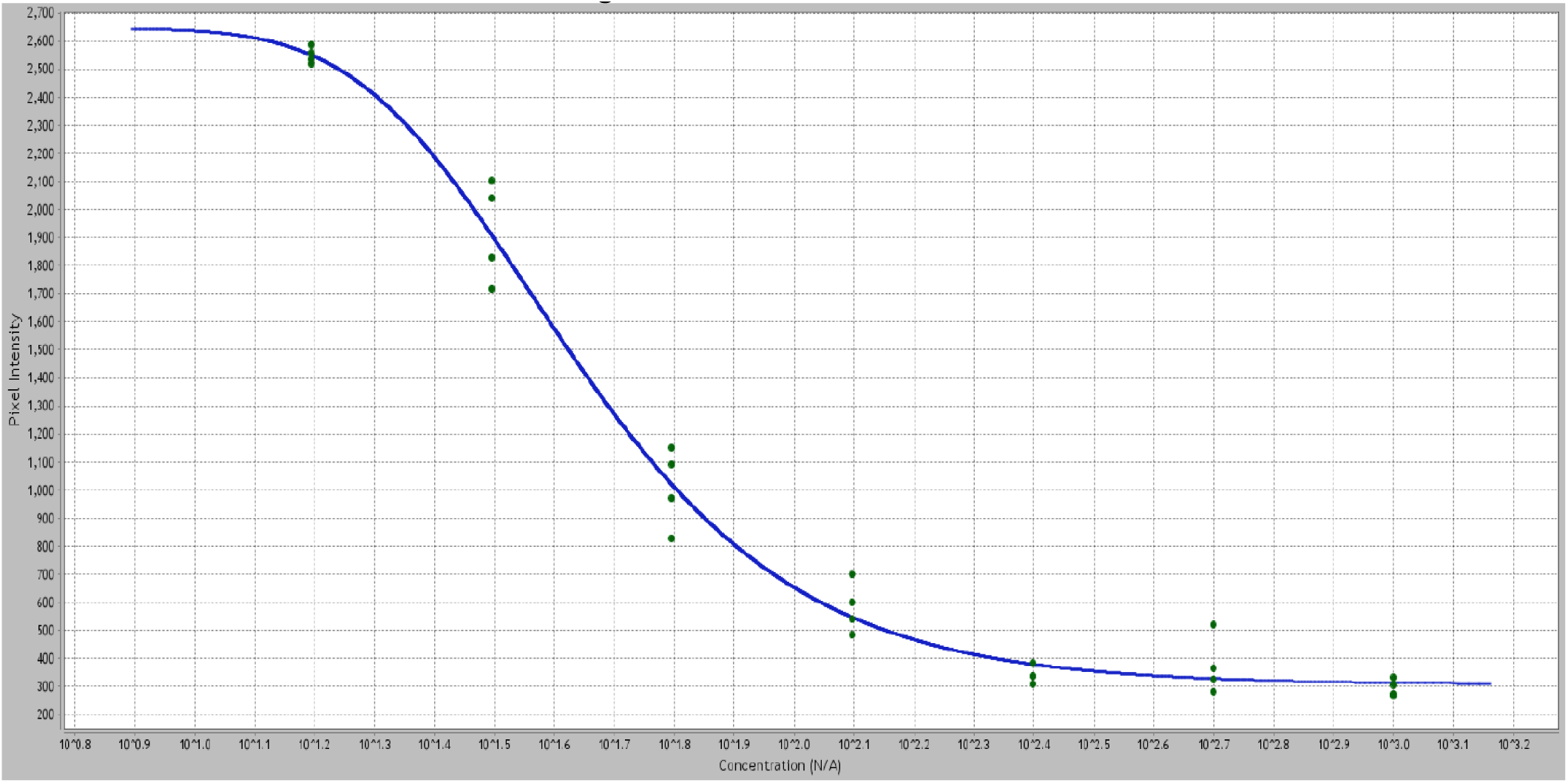
Standard Curve for FOLR1-Folate Binding. Signals are extracted and plotted as pixel intensity (Y) versus folic acid concentration (X) to produce a standard FOLR-folate binding curve. Increased concentration (60 - 0.9 ng/mL, dilutions in quadruplicate) of unlabeled folic acid in the presence of a constant FA-HRP results in decreased signal intensity.

**Supplemental Figure 2.**
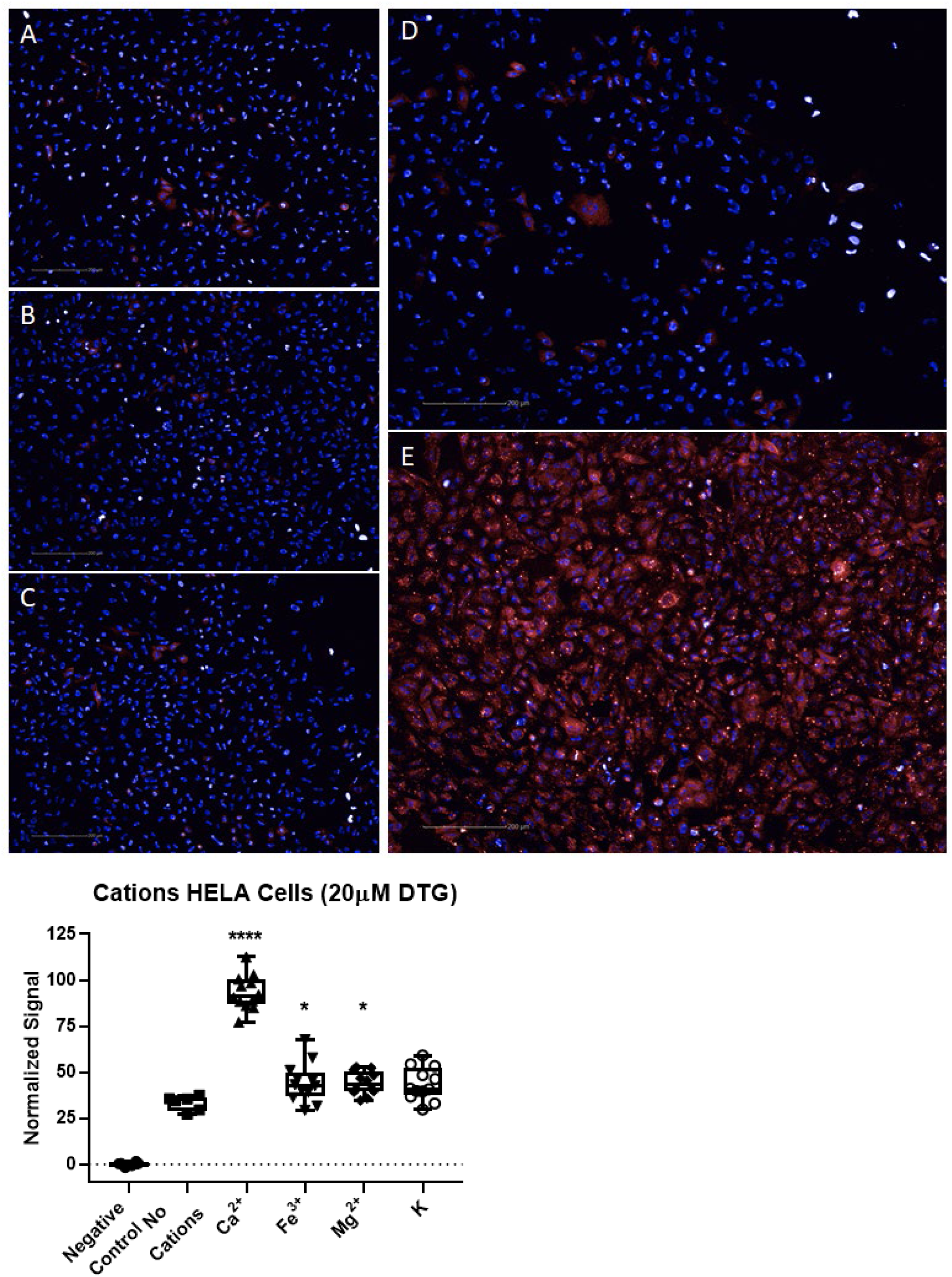
Calcium Modifies Cellular Folate Binding. The impact of cations on folate binding in the presence of DTG was initially tested in HELA cells. Images of HELA cells exposed to 20μM DTG, 50nM FA-680 (red), and Hoechst 3342 (blue, live-nuclei) in buffer (1X DPBS, 1% FBS). The impact of A) potassium, B) magnesium, C) iron, D) control, and E) calcium are shown at 24hrs. Potassium showed a non-significant increase in FA-680 signal. Iron and magnesium showed significant increased FA-680 signal (* P<0.05). Calcium showed the largest increase in cellular FA-680 binding compared to control (**** P<0.0001).

**Supplemental Figure 3.**
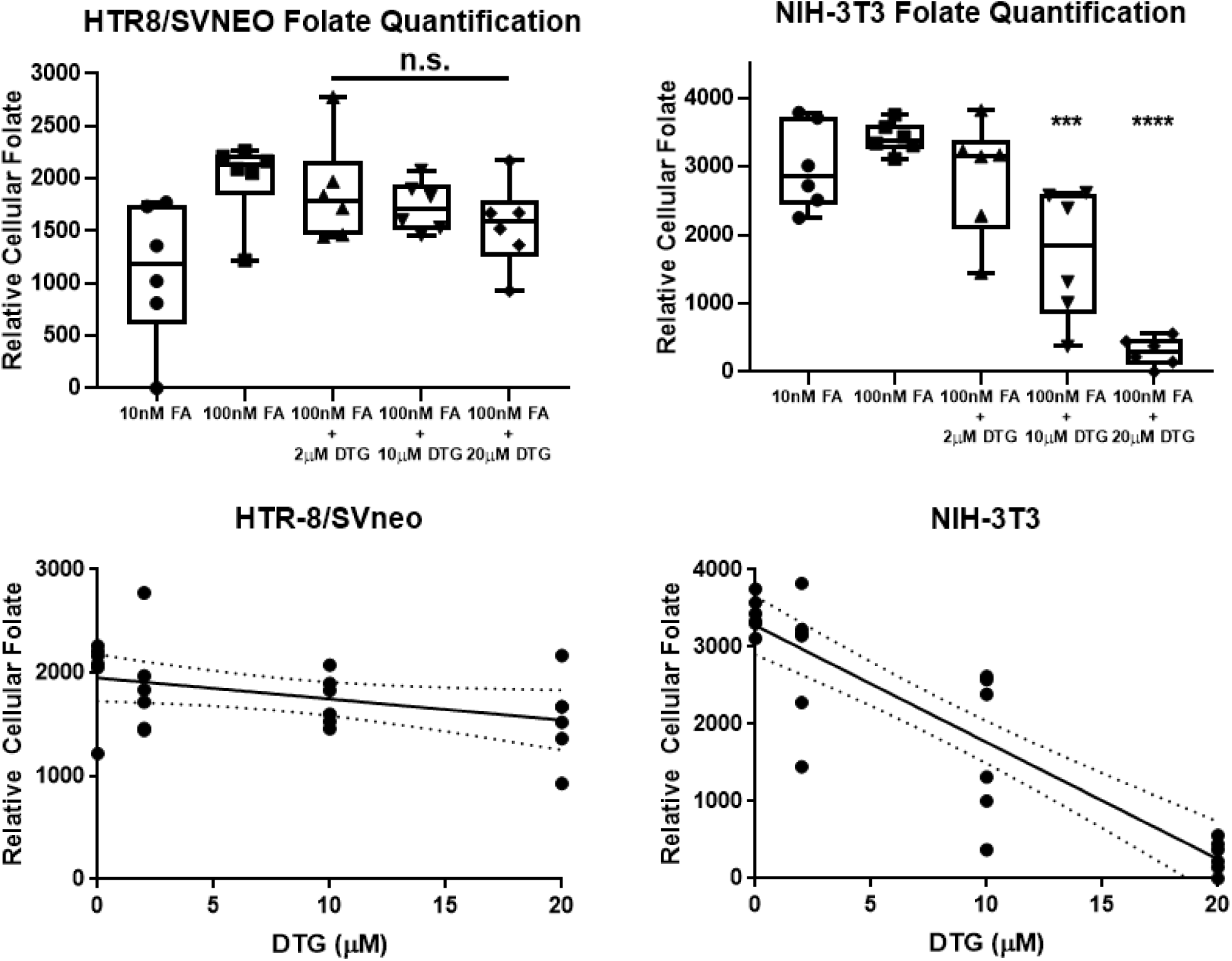
Impact of DTG on Folate Uptake in HTR-8 and NIH3T3. The HTR8/SVneo were cultured in RPMI (without folate) and NIH-3T3 were cultured in DMEM (without folate) and various exposures were tested for interactions in folate (unlabeled) uptake. The folate concentration in cell pellets were determined by FOLR1 protein binding assays and normalized to total cellular protein. There was non-significant decreased uptake of folate by DTG (2-20μM) in HTR-8 cells. In NIH-3T3 mouse fibroblasts, there was a significant decrease (*** P < 0.001, **** P <0.0001) in folate uptake at 10 μM and 20μM DTG. The NIH-3T3 are cultured in media (DMEM) that contains 1.8mM Ca^+^2, and the HTR8/SVneo contains 0.42mM Ca^+^2.

